# BioLab: End-to-End Autonomous Life Sciences Research with Multi-Agents System Integrating Biological Foundation Models

**DOI:** 10.1101/2025.09.03.674085

**Authors:** Ruofan Jin, Yucheng Guo, Yuanhao Qu, Ming Yang, Chun Shang, Qirong Yang, Linlin Chao, Yi Zhou, Ruilai Xu, Ziyao Xu, Ruhong Zhou, Zaixi Zhang, Mengdi Wang, Xiaoming Zhang, Le Cong

**Author notes:** **Additional information Correspondence and requests for materials** should be addressed to Ruhong Zhou, Zaixi Zhang, Mengdi Wang, Xiaoming Zhang, and Le Cong., and. Equal Contribution (Ruofan Jin, Yucheng Guo, Yuanhao Qu).

## Abstract

Scientific discovery in the life sciences remains hindered by fragmented workflows, narrow-scope computational models, and inefficient links between in silico prediction and wet-lab validation. We present BioLab, a multi-agent system that integrates domain-specialized foundation models to automate end-to-end biological research. BioLab comprises eight collaborating agents, including a Planner, Reasoner, and Critic, orchestrated through a Memory Agent that enables iterative refinement via retrieval-augmented generation and a suite of 219 computational xBio-Tools spanning five biological scales (DNA, RNA, protein, cell, and chemical). These tools are built on the xTrimo Universe, a collection of 104 models derived from six foundation models (xTrimoChem, Protein, RNA, DNA, Cell, and Text), the majority of which achieve state-of-the-art (91–100% SOTA ratios) on domain benchmarks. Across standard reasoning tasks (PubMedQA, MMLU-Pro/Biology, GPQA-diamond), BioLab consistently outperformed leading large language models, including GPT-4o, Gemini-2.5, and DeepSeek-R1. Beyond benchmarks, BioLab autonomously executed a fully computational pipeline for de novo macrophage-targeting antibody design, progressing from target mining to multi-objective antibody optimization, where molecular dynamics simulations revealed structural mechanisms underlying enhanced affinity of optimized variants. Closing the computational-experimental loop, BioLab designed optimized antibodies (Pem-MOO-1, Pem-MOO-2) that achieved IC50 values of 0.01–0.016 nM, markedly surpassing the parental Pembrolizumab (0.027 nM) for PD-1. Functional assays confirmed enhanced pathway blockade and improved multi-parameter performance profiles. Together, these results establish BioLab as a generalizable framework for AI-native scientific discovery, demonstrating how multi-agent systems coupled with foundation models can autonomously generate, execute, and experimentally validate novel biological hypotheses.

## Introduction

Scientific discovery in the life sciences is increasingly defined by its scale and complexity^1, 2^, demanding the integration of multimodal data and iterative experimentation across disciplines^3^. From genomics and structural biology to systems-level immunology, modern investigations generate vast datasets at an unprecedented rate^4^. Yet, despite these advances in data acquisition, the scientific workflow itself remains deeply fragmented^5^. Biological knowledge is siloed in specialized subfields^6, 7^, analytical tools are often non-interoperable^8^, and the critical feedback loop between computational modeling and experimental validation is inefficient and laborious^9, 10^. These discontinuities create a significant bottleneck, impeding hypothesis generation, leading to missed opportunities for synergistic insights, and fundamentally limiting the pace of discovery^11^.

Foundation models have emerged as powerful instruments in computational biology, enabling significant advances in structure prediction, sequence design, and property inference^12–16^. These models, however, are typically applied as isolated point solutions, optimized for singular tasks or data modalities^17–19^. They lack the executive function required to navigate the full arc of scientific inquiry from conceptual reasoning and strategic planning to experimental design and data interpretation^19, 20^.

For example, they cannot independently formulate a multi-stage research hypothesis^21^, allocate computational resources^22^, or interpret the results of one experiment to design the next^12, 23^. Consequently, while individual tasks have been accelerated, the broader vision of an integrated, model-driven research system remains unrealized.

General-purpose large language models (LLMs) have introduced new paradigms for multi-step reasoning and agentic task execution^24^. While these systems hint at a future of AI-augmented science, they are not inherently fluent in the language of biology^25^. Their capabilities are often limited to superficial linguistic fluency, which is insufficient for the deep, causal, and mechanistic understanding that biological research demands^26, 27^. Lacking deep domain-specific grounding, they are prone to hallucination and cannot reliably engage with the rich, specialized ecosystem of computational models, databases, and experimental protocols that form the bedrock of the field^28–30^. In a domain where precision and fidelity are paramount, their generalist nature becomes a critical liability^28, 31^.

To address these challenges, we developed **BioLab**, a multi-agent system purpose-built to automate and unify the end-to-end life sciences research workflow. BioLab orchestrates a synergistic community of modular expert agents, a retrieval-augmented knowledge base, and a toolkit of specialized computational models derived from a core set of foundation models into a single, coherent framework. Its agents coordinate to propose hypotheses, query biomedical databases, invoke predictive models, and, critically, generate actionable experimental protocols and incorporate the results into subsequent rounds of inquiry. We demonstrate BioLab’s capabilities through two progressively challenging case studies. First, we show its capacity for end-to-end automation in a fully in silico workflow to identify macrophage-specific targets and engineer a corresponding antibody. Second, and more stringently, we establish a closed-loop integration with the wet lab, where BioLab guides the discovery of a T-cell target and optimizes a therapeutic antibody, leading to predictions that are prospectively validated by physical experiments. These results establish BioLab as a robust framework for AI-native scientific discovery, providing a blueprint for the future of automated, hypothesis-driven research in biology.

### BioLab: A multi-agent AI system to automate the scientific discovery lifecycle

A fundamental challenge in modern life sciences is the fragmentation of the research process. Biological knowledge is siloed across specialized subdomains, computational models are often confined to narrow, isolated tasks, and the critical feedback loop between experimental results and subsequent reasoning is slow and inefficient^32, 33^. To overcome these barriers, we developed BioLab, a virtual laboratory powered by a modular multi-agent AI system designed to autonomously orchestrate the entire scientific discovery lifecycle (Figure 1a). The architecture is engineered around the core principles of division of labor, memory continuity, and scientific reliability. A key innovation of BioLab is its ability to understand and navigate the complexities of the research process, seamlessly integrating multi-round interactions between computational modeling and wet-lab experimentation. This capability allows BioLab to automate the full “design-build-test-learn” cycle, establishing a truly closed-loop system that iteratively refines hypotheses based on emergent evidence.

**Figure 1.**
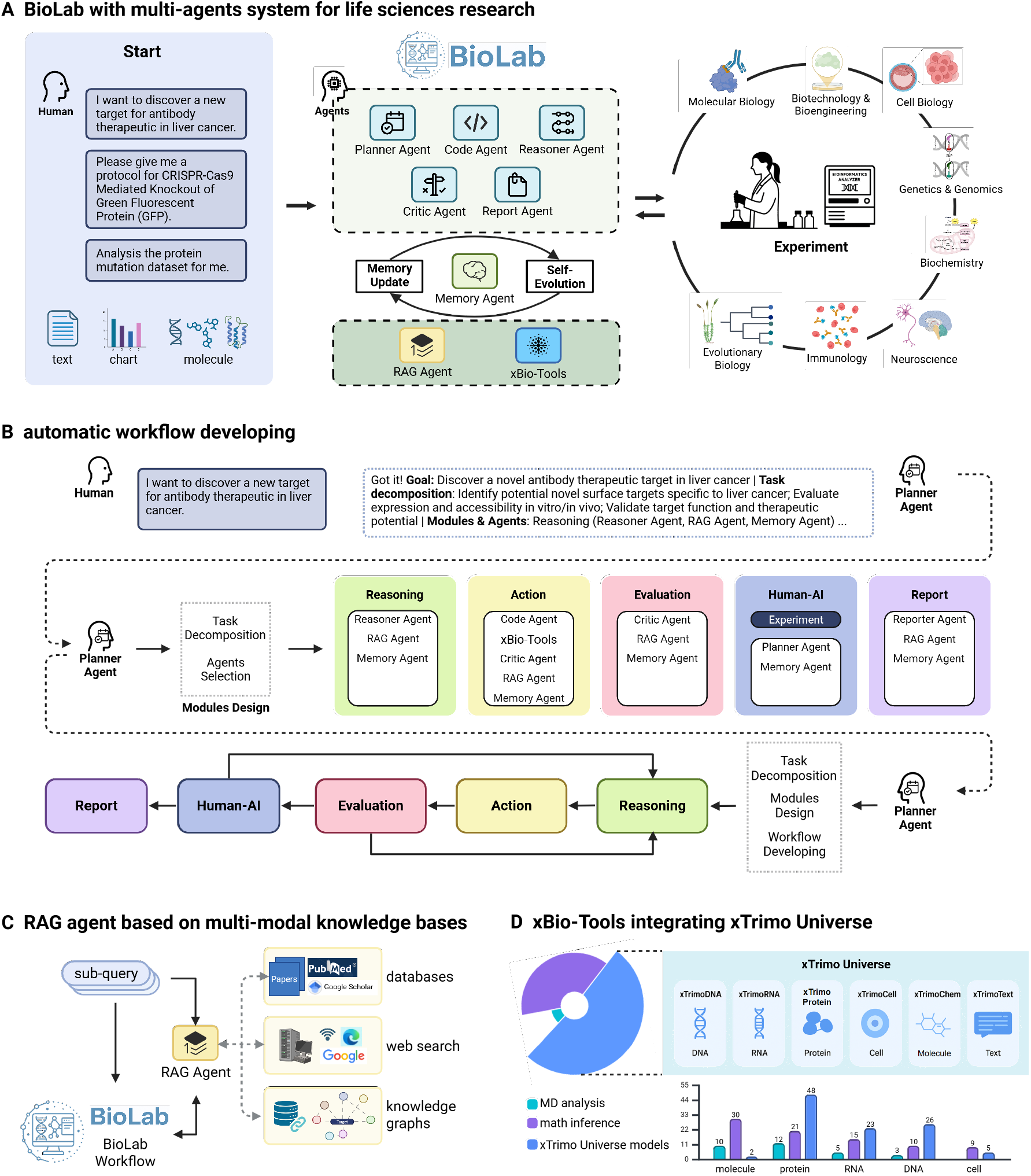
BioLab, a multi-agent system, orchestrates foundation models for automated scientific discovery. (**A**) The architecture of BioLab. User instructions, including natural language queries and multimodal inputs (text, charts, molecular data), are processed by a multi-agent system. The system comprises eight core agents (Planner, Reasoner, Critic, etc.) that collaborate to execute complex tasks. A Memory Agent enables self-evolution by updating the knowledge base used by the RAG Agent and xBio-Tools, creating a closed loop between in silico prediction and wet-lab experimental design. (**B**) Automated workflow generation. The Planner Agent decomposes a user’s primary objective into discrete tasks. It then dynamically assembles functional modules (e.g., Reasoning, Action, Evaluation) and sequences them into an optimal workflow. This process is iterative, allowing for continuous refinement based on intermediate results. (**C**) Retrieval-Augmented Generation (RAG) for knowledge synthesis. Sub-queries derived from the main task are addressed by the RAG Agent. It synthesizes information from diverse sources, including literature databases (PubMed, Google Scholar) (Extended Data Figure S1a), web searches (Extended Data Figure S1a), and biomedical knowledge graphs (Extended Data Figure S1a), to provide comprehensive, context-aware answers (Extended Data Figure S1b). (**D**) xBio-Tools powered by the xTrimo Universe. The system integrates 219 computational tools for analyses across five biological scales (DNA, RNA, protein, cell, chemical). These tools are built upon the xTrimo Universe, a suite of 104 models derived from six foundational models, each specialized for a specific data modality. Bar chart quantifies the number of tools available for each biological scale. See Extended Data Table for the complete xBio-Tools list.

The operational core of BioLab is its autonomous workflow generation, orchestrated by a dedicated Planner Agent (Figure 1b). Upon receiving a high-level user query, the Planner Agent algorithmically deconstructs the objective into a logical sequence of subtasks. It then assembles the requisite agents and computational tools into functional modules (e.g., Reasoning, Action, Evaluation) and dynamically sequences them into a coherent, multi-step workflow. This process is underpinned by a Memory Agent (see Memory Agent section in Methods for details.) that captures structured representations of all intermediate findings, including external experimental data. This persistent memory is crucial for longitudinal learning, allowing BioLab to maintain context and adapt its strategy across multiple rounds of inquiry, mirroring the iterative process of human scientific reasoning but at a vastly accelerated pace.

To ensure that all reasoning is grounded in robust scientific knowledge, BioLab employs a Retrieval-Augmented Generation (RAG) agent operating on a bespoke, multimodal biomedical knowledge base (Figure 1c). This knowledge base was custom-built to overcome the limitations of relying on a single information source. It integrates three distinct, synergistic components: (i) a comprehensive corpus of scientific literature for foundational knowledge; (ii) real-time web-retrieval capabilities to access the most current, rapidly evolving information; and (iii) a proprietary, large-scale biomedical knowledge graph that explicitly models the complex relationships between genes, proteins, diseases, and drugs. This tripartite architecture, powered by a domain-specific Text2NGQL model we developed (Extended Data Figure S1), facilitates a graph-based RAG process that substantially improves factual grounding and reduces the incidence of hallucination compared to standard methodologies (Extended Data Table S3).

The execution of specific biochemical and biophysical analyses is handled by xBio-Tools, a comprehensive suite of 219 validated computational tools (Figure 1d). This toolkit spans five fundamental biological scales—from chemical structures to cellular systems—and is powered by the xTrimo Universe, a collection of 104 predictive models we developed from six large-scale foundation models (see the xTrimo Foundation Models section in Materials and Methods for details). These models, trained on vast biological datasets, provide domain-aware priors for downstream reasoning and can be invoked for high-fidelity tasks such as structure prediction and biophysical simulation. Collectively, BioLab’s architecture integrates strategic planning, deep-domain knowledge, and high-performance computing into a single, unified platform for automated scientific discovery.

### BioLab demonstrates superior performance on core scientific reasoning and life science modeling tasks

To validate the core cognitive abilities required to overcome research fragmentation, we quantitatively evaluated BioLab’s performance against existing state-of-the-art (SOTA) AI systems. We first assessed its capacity for deep scientific reasoning on a panel of four challenging biomedical question-answering benchmarks (PubMedQA (with and without content)^34^, MMLU-Pro/Biology^35^, and GPQA-diamond^36^), which are designed to test comprehension of nuanced and highly specialized knowledge. Across all benchmarks, BioLab consistently outperformed leading large language models (LLMs), including DeepSeek-R1^37^, Qwen3-235B-A22B^38^, ChatGPT-4o^39, 40^ and Gemini-2.5-flash^41, 42^, underscoring its superior ability to reason within the complex domain of biology (Figure 2a).

**Figure 2.**
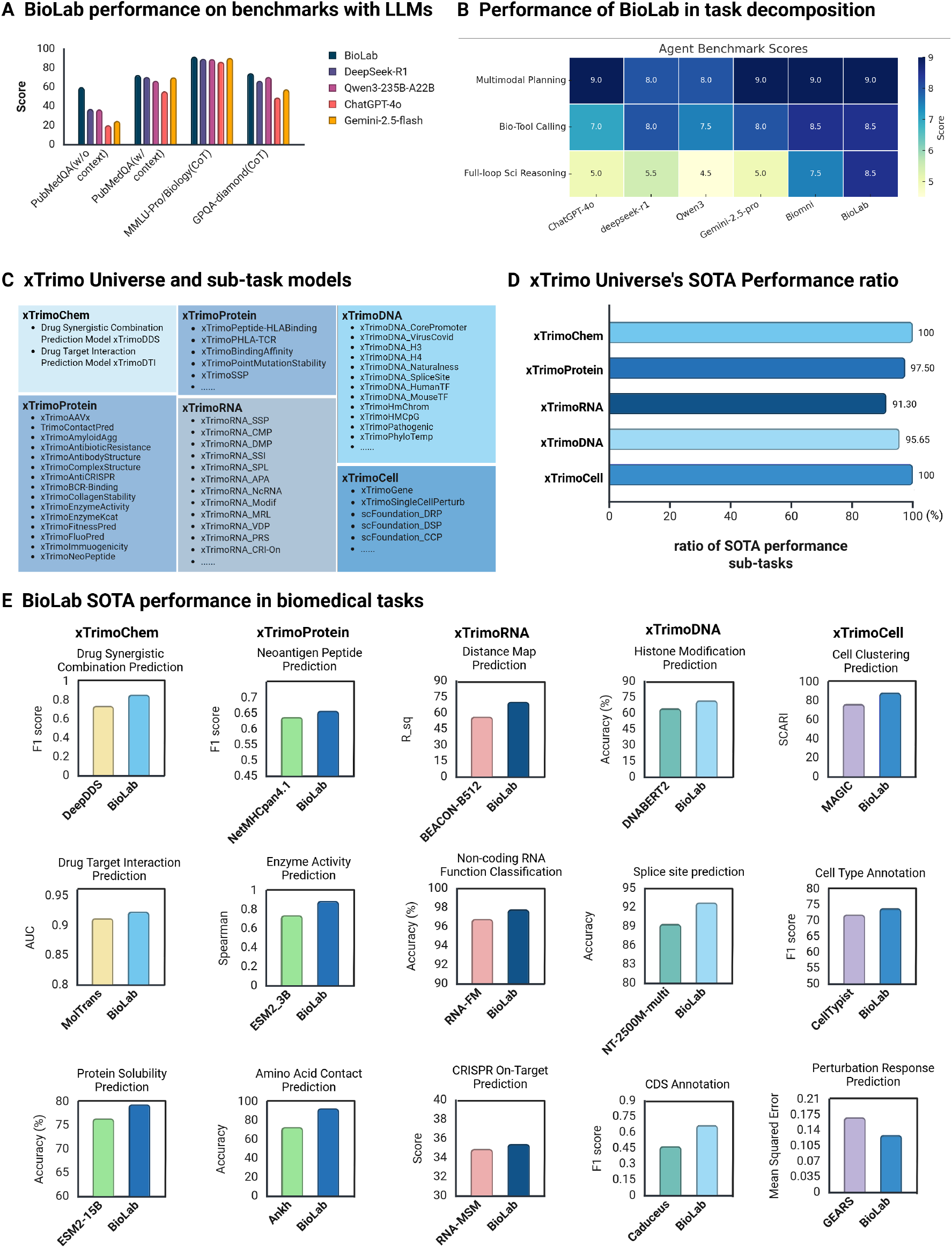
Performance benchmarks and state-of-the-art capabilities of BioLab and its core xTrimo Universe. (**A**) Superior reasoning performance on biomedical benchmarks. BioLab’s performance was evaluated against leading large language models (LLMs), including DeepSeek-R1, Qwen3-235B, OpenAI-GPT-4o, and Gemini-2.5-flash, on four scientific question-answering benchmarks: PubMedQA (with and without content), MMLU-Pro/Biology (CoT), and GPQA-diamond (CoT). BioLab consistently achieved higher scores across all tested benchmarks. (**B**) State-of-the-art performance in automated task decomposition. On the BioResearchQA benchmark, BioLab’s ability to decompose complex tasks was compared against other AI models using three expert-defined evaluation metrics. BioLab demonstrated superior performance across all metrics, outperforming both LLMs and the Biomni agent model. (**C**) The xTrimo Universe of foundation and task-specific models. The xTrimo Universe comprises six foundation models, each targeting a different biological scale (xTrimoChem, xTrimoProtein, xTrimoRNA, xTrimoDNA, xTrimoCell, and xTrimoText), along with a comprehensive suite of downstream task-specific models. A representative subset of the model collection is shown. See Extended Data Table for the complete list. (**D**) Quantification of state-of-the-art (SOTA) models within the xTrimo Universe. The bar chart illustrates the proportion of downstream models within each category of the xTrimo Universe that achieve SOTA performance on their respective domain-specific benchmarks. The results indicate a high prevalence of SOTA models, with ratios ranging from 91.30% to 100%. (**E**) SOTA performance on representative biomedical tasks. Bar plots show the SOTA performance of selected xTrimo downstream models on a variety of critical biomedical tasks, including drug synergy prediction (DeepDDS), enzyme activity prediction (ESM2-3B), splice site prediction (NT-2500M-multi), and cell type annotation (CellTypist). See Extended Data Table S4 for a comprehensive evaluation of all downstream models.

A cornerstone of automating research is the ability to deconstruct a high-level goal into a logical, multi-step experimental plan^43^. We therefore evaluated BioLab’s performance on the BioResearchQA benchmark, which specifically assesses this task-decomposition capability. Using three independent, expert-defined metrics that probe different aspects of planning, BioLab again demonstrated superior performance, outperforming both general-purpose LLMs and other specialized agent-based models (Figure 2b). This result highlights the effectiveness of our Planner Agent in formulating scientifically valid and logically coherent research strategies, a critical step in orchestrating complex, end-to-end projects (see the Benchmarking and evaluation methodology section in Methods for details).

The practical utility of BioLab’s plans depends on the quality of the tools it orchestrates. The system’s power is derived in large part from its integrated xTrimo Universe, a vast collection of foundation and task-specific models (Figure 2c). We systematically benchmarked our suite of downstream models against established SOTA models in their respective domains. The results revealed an exceptionally high prevalence of SOTA-level performance, with the proportion of our models achieving this status ranging from 91.30% to a full 100% in key categories like xTrimoChem and xTrimoCell (Figure 2d).

To provide concrete evidence of these capabilities, we highlight several examples where xTrimo models excel in critical biomedical tasks that are fundamental to drug discovery and mechanistic biology (Figure 2e). For instance, in predicting drug synergy, modeling protein interactions, mapping RNA accessibility, and annotating cell types, our models consistently matched or surpassed the performance of previous benchmark holders. This integrated suite of high-performing, specialized tools is what allows BioLab to move beyond purely linguistic planning to perform high-fidelity, quantitative biological modeling, thereby executing the complex workflows it designs. Complete results on BioLab’s SOTA performance are provided in Extended Data Table S4.

### BioLab executes a closed-loop in silico workflow for macrophage target discovery and antibody engineering

To demonstrate BioLab’s capacity to orchestrate a complete research cycle in silico, we tasked it with a complex, real-world challenge that typically requires months of human effort: designing a therapeutic antibody targeting macrophages for cancer treatment. Upon receiving this single, high-level command, BioLab, without any further human intervention, autonomously designed and executed a five-stage discovery workflow (Figure 3a). This cascade of intelligent actions seamlessly integrated target discovery, computational validation, antibody sourcing, and multi-objective antibody optimization, showcasing a fully automated, closed-loop research project.

**Figure 3.**
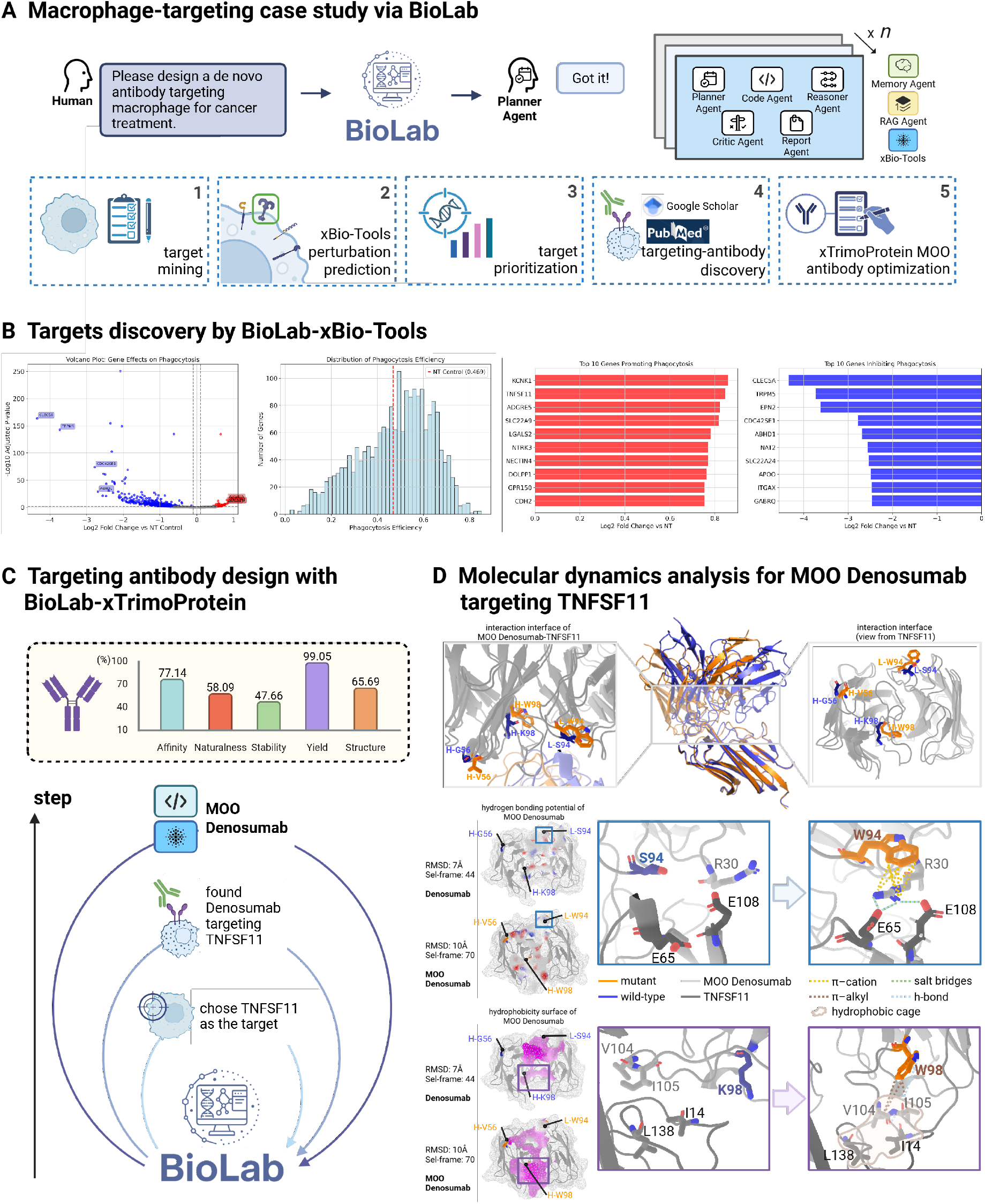
End-to-end de novo design and optimization of a macrophage-targeting antibody by BioLab. (**A**) An automated, end-to-end workflow for macrophage-targeting antibody discovery. Upon receiving a high-level user command (’Please design a de novo antibody targeting macrophage for cancer treatment’), the Planner Agent autonomously orchestrates a multi-stage workflow. This process includes five key phases: (1) target mining: target mining from literature and databases; (2) xBio-Tools perturbation prediction: target discovery and screening via xBio-Tools perturbation prediction; (3) target prioritization: prioritization of candidate targets; (4) targeting antibody discovery: discovery of existing antibodies against high-priority targets; and (5) xTrimoProtein MOO antibody optimization: multi-objective optimization (MOO) of a selected antibody using the xTrimoProtein model. This entire pipeline operates as a closed-loop, automated system. (**B**) Identification of potential macrophage targets using xBio-Tools. BioLab identifies candidate targets by analyzing transcriptomic data. The volcano plot (left) highlights differentially expressed genes, pathway enrichment analysis (center) identifies relevant biological processes, and the bar charts (right) rank top candidate targets based on expression levels and biological significance. Notably, TNFSF11 was identified among the top-ranked candidates. (**C**) Multi-objective optimization of the anti-TNFSF11 antibody, Denosumab. BioLab identified TNFSF11 as a high-priority target and retrieved the corresponding antibody, Denosumab, from its knowledge base. Subsequently, the system employed the xTrimoAntibody model, a downstream application of xTrimoProtein, to perform MOO. The optimization process simultaneously enhanced five key properties: affinity, naturalness, stability, yield, and structural integrity, resulting in an improved variant, MOO Denosumab. (**D**) Molecular dynamics (MD) simulations reveal the structural basis for enhanced affinity. MD simulations elucidated the mechanism underlying the improved binding of MOO Denosumab. The optimized antibody (orange) features three key mutations (H-G56V, H-K98W, L-S94W) compared to the wild-type (WT) structure (blue). The L-S94W mutation induces a conformational change via a *π*–cation interaction (yellow dashed line) with R30, which in turn forms a new hydrogen bond (blue dashed line) and a salt bridge (green dashed line) with residues E65 and E108 on the antigen. In concert, the H-K98W mutation introduces additional hydrophobic interactions (including *π*–alkyl contacts; brown dashed line) that organize surrounding residues into a stable hydrophobic cage (tan surface), further reinforcing the antibody-antigen interface.

The process began with target identification, a critical and often laborious phase of drug discovery. BioLab initiated a systematic search, mining public databases and the scientific literature to identify genes differentially expressed in tumor-associated macrophages. It then leveraged the xTrimoSingleCellPerturb model within xBio-Tools suite to perform in silico perturbation predictions and pathway analyses on the initial hits, ultimately prioritizing a list of candidate targets based on their predicted functional impact (Figure 3b). Notably, TNFSF11^44–47^, a well-established signaling protein and the target of the clinical antibody Denosumab^48–51^, was identified among the top-ranked candidates, validating the system’s ability to independently pinpoint biologically and therapeutically relevant targets (see the xTrimo platform integration and foundation model usage within BioLab section in Methods for details).

Once TNFSF11 was confirmed as the primary target, BioLab transitioned to the antibody engineering phase. It first queried its knowledge base to retrieve the sequence and structural information for Denosumab (see the BioLab-Guided Antibody Recommendation with Bioinformation for TNFSF11 section in Materials and Methods for details). The system then employed the xTrimoAntibody model to formulate and execute a multi-objective optimization (MOO) on the Denosumab sequence—a computationally demanding task that requires navigating a high-dimensional search space to balance multiple, often conflicting, biophysical parameters (Figure 3c). The goal is to simultaneously improve five key properties critical for therapeutic success: binding affinity, naturalness, stability, production yield, and overall structural integrity.

The MOO process yielded a novel, optimized variant, designated MOO Denosumab (Figure 3c), with a predicted property profile superior to the wild-type antibody. This demonstrates BioLab’s ability to not only identify existing therapeutic candidates but to rationally engineer them for enhanced performance. The system’s capacity to handle such complex, multi-parameter optimization problems is a crucial step towards AI-driven molecular design.

To provide a mechanistic explanation for the predicted improvements–—a hallmark of deep scientific inquiry—–we directed BioLab to conduct molecular dynamics (MD) simulations^52, 53^ on both the wild-type (WT) and the MOO Denosumab–TNFSF11 complexes (Figure 3d). The simulations revealed that the three mutations introduced during optimization induced critical, cooperative conformational changes at the binding interface. Specifically, the L-S94W mutation facilitated a new *π*–cation interaction that stabilized the formation of a new hydrogen bond and salt bridge, while the H-K98W mutation established novel hydrophobic interactions (including *π*-alkyl contacts) that organized a more stable hydrophobic cage. These atomic-level insights not only explain the enhanced binding affinity but also demonstrate BioLab’s ability to generate testable, mechanistic hypotheses, moving far beyond simple prediction to offer deep biological understanding.

### BioLab closes the computational-experimental loop for T-cell target discovery and anti-body optimization

The ultimate test of any computational discovery platform is its ability to generate novel, non-obvious predictions that are subsequently validated by real-world experiments. To this end, we designed a final study to prospectively validate BioLab’s capabilities in a complete wet-lab/dry-lab cycle, tasking it with discovering a therapeutic strategy for T-cell-based cancer immunotherapy. We established a fully integrated,”design-build-test-learn” cycle, where BioLab’s computational predictions were directly followed by experimental validation, with the results feeding back into the system to inform the next steps (Figure 4a). This case study was designed to rigorously test BioLab’s ability to bridge the gap between in silico hypothesis and tangible, experimentally verified outcomes.

**Figure 4.**
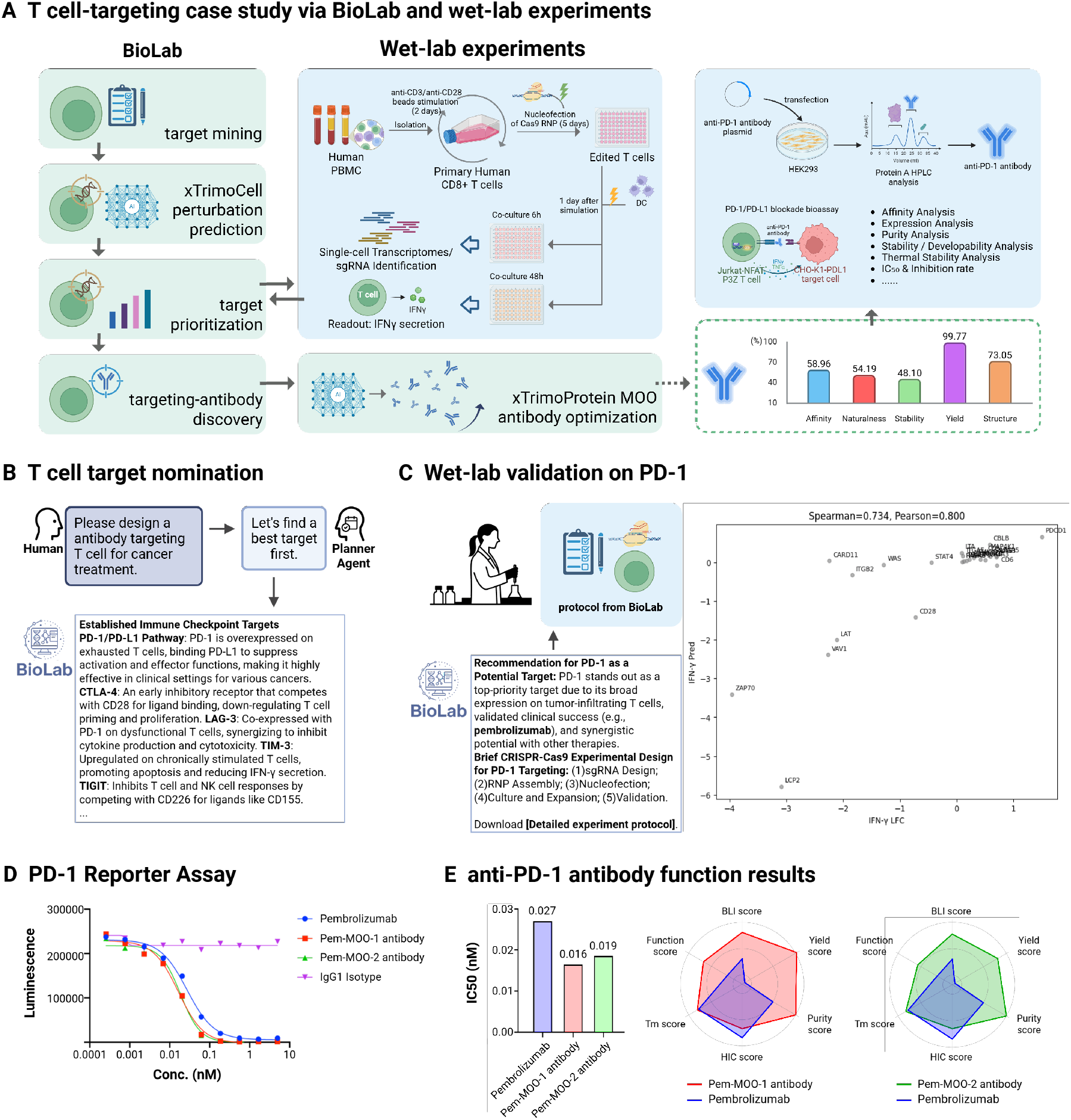
Closed-loop integration of BioLab and wet-lab experiments for T-cell target discovery and antibody optimization. (**A**) A hybrid computational/wet-lab workflow for T-cell therapy development. BioLab (in silico, green boxes) and wet-lab experiments (blue boxes) were integrated into a multi-round “design-build-test-learn” cycle. The in silico pipeline includes target mining, xTrimoCell-based perturbation prediction, target prioritization, antibody discovery, and xTrimoProtein-driven antibody optimization. The wet-lab pipeline involves isolating primary human CD8+ T cells from PBMCs, performing CRISPR-Cas9 gene editing, co-culturing with dendritic cells (DCs) for IFN*γ* secretion assays, and validating antibody function in a HEK293-based PD-1/PD-L1 blockade bioassay. (**B**) Autonomous identification of PD-1 as the top T-cell target. Upon receiving the high-level command ’design an antibody targeting T cell for cancer treatment’, BioLab’s reasoning engine analyzed established immune checkpoint pathways and prioritized Programmed Death-1 (PD-1) as the lead candidate with the highest therapeutic potential. (**C**) Wet-lab validation confirms that PD-1 knockout enhances T-cell activity. BioLab generated a detailed protocol for CRISPR-Cas9-mediated knockout of PDCD1 (the gene encoding PD-1), integrating multi-agent analysis and the xTrimoSingleCellPerturbation model to prioritize PD-1. Subsequent experimental data revealed a strong positive correlation (Spearman correlation: 0.734, Pearson correlation: 0.800) between the model-predicted outcomes and the wet-lab results of IFN*γ* secretion score. This result functionally validates BioLab’s target recommendation by directly linking reduced PD-1 expression to enhanced T-cell immune activity, aligning computational predictions with experimental outcomes. (**D**) Performance of optimized antibodies in a PD-1/PD-L1 blockade reporter assay. The ability of the antibodies to block the PD-1/PD-L1 interaction was assessed in a HEK293-based reporter assay. The dose-response curves show that both multi-objective optimized (MOO) variants, Pem-MOO-1 and Pem-MOO-2, exhibited more potent pathway blockade than the parental antibody, Pembrolizumab, indicated by a stronger restoration of the luminescence signal at lower concentrations. The IgG1 isotype control showed no activity. (**E**) Comprehensive functional and biophysical characterization of optimized antibodies. Quantification of IC50 values (left) demonstrates that the optimized variants, Pem-MOO-1 (0.016 nM) and Pem-MOO-2 (0.01 nM), are substantially more potent than the parental antibody, Pembrolizumab (0.027 nM). The radar plots (right) provide a multi-parameter assessment of both functional attributes (e.g., BLI score, Function score) and biophysical properties (e.g., Purity, Yield), with all metrics scaled so that a larger area indicates a more desirable profile. The profile for Pem-MOO-2 reveals substantial improvements in key functional metrics—notably the Function and BLI scores—while maintaining comparable performance in biophysical properties such as yield and purity. This enhanced efficacy was achieved with an acceptable trade-off in hydrophobicity (HIC score). The significantly larger polygonal area of the Pem-MOO-2 profile underscores a superior overall candidate, validating the success of the multi-objective optimization strategy in substantially boosting antibody function while managing biophysical liabilities.

Given the high-level goal, BioLab’s reasoning engine first analyzed established immune checkpoint pathways and prioritized Programmed Death-1 (PD-1) as the lead candidate target. This step alone replicates a complex decision-making process that typically involves extensive literature review and expert consultation (Figure 4b). This demonstrates the system’s ability to distill vast amounts of information into a single, high-confidence starting point for a research campaign, effectively overcoming the initial hurdle of hypothesis generation.

Crucially, to bridge the gap from strategy to execution, BioLab then generated a complete, step-by-step laboratory protocol for knocking out the PDCD1 gene (which encodes PD-1) in primary human T-cells using CRISPR-Cas9 technology^54–56^. The ability to generate such actionable, detailed experimental plans is a major advance toward true research automation, as it directly addresses the practical barrier between computational hypothesis and experimental action. The generated protocol provided specific reagents, concentrations, and incubation times required for the experiment.

The wet-lab experiments were conducted following the BioLab-generated protocol (see Materials and Methods), seamlessly extending the computational framework into experimental validation. Building on BioLab’s multi-agent system, which leveraged literature and knowledge to prioritize PD-1 as a key T-cell target, and its use of the xTrimoSingleCellPerturb model within xTrimo Universe to rank PD-1 first among predicted gene perturbations, the study incorporated parallel wet-lab efforts. These experiments confirmed PD-1’s top ranking through T-cell gene perturbation assays based on IFN-predictions, revealing a strong positive correlation between reduced PD-1 protein expression and enhanced T-cell immune activity (Spearman correlation: 0.734, Pearson correlation: 0.800; Figure 4c). This robust alignment between computational forecasts and experimental outcomes not only validates BioLab’s predictive accuracy but also reinforces its role in guiding the development of effective T-cell-based therapeutic strategies, paving the way for subsequent antibody optimization.

With the target validated, we leveraged BioLab to complete the therapeutic discovery cycle by optimizing Pembrolizumab, a clinical antibody targeting PD-1 (see Materials and Methods). The system generated two optimized variants, Pem-MOO-1 and Pem-MOO-2, which were then synthesized and tested in a functional reporter assay. Both variants exhibited more potent pathway blockade than the parental antibody in a dose-dependent manner. Notably, Pem-MOO-2 showed a nearly three-fold improvement in IC50 (0.01 nM vs. 0.027 nM for parental), demonstrating a substantial enhancement in biological activity resulting directly from the in silico optimization campaign (Figure 4d and Figure 4e, left).

Finally, a comprehensive multi-parameter assessment of the optimized antibodies was performed to evaluate their overall therapeutic potential and ensure that functional gains did not come at the cost of developability. A normalized radar plot revealed that Pem-MOO-2 not only possessed superior potency and binding affinity (Function and BLI scores) but also maintained developability profiles, such as yield, purity, and thermostability, comparable to the parental antibody (Figure 4e, right). This enhanced efficacy was achieved with an acceptable and well-understood trade-off in hydrophobicity (HIC score), a common challenge in antibody engineering. The significantly larger polygonal area of the Pem-MOO-2 profile underscores a superior overall candidate, proving that BioLab can successfully navigate the complex, multi-parameter landscape of therapeutic optimization. This final result successfully closes the loop, demonstrating that BioLab can shorten the discovery cycle by autonomously navigating from high-level strategy to the engineering of superior therapeutic molecules with experimentally confirmed efficacy.

## Discussion

Here we present BioLab, a multi-agent AI system designed to address the profound fragmentation of the modern life sciences research workflow. We have demonstrated that by orchestrating a synergistic community of specialized agents, BioLab can automate the entire scientific discovery lifecycle, from high-level hypothesis generation to the design of superior therapeutic molecules with experimentally validated efficacy. Our results establish a robust framework for a new paradigm of AI-native scientific research, capable of closing the loop between computational modeling and physical experimentation.

The core innovation of BioLab lies in its architecture, which overcomes the limitations of both isolated foundation models and general-purpose LLMs. While foundation models have revolutionized specific tasks such as protein structure prediction, they function as specialized tools and lack the executive function to orchestrate a multi-stage research campaign^21, 57, 58^. Conversely, generalist LLMs, while capable of reasoning, lack the deep domain grounding necessary for high-fidelity biological inquiry, making them prone to factual hallucination^30, 59, 60^. BioLab resolves this dichotomy by integrating a hierarchical agent system, where a Planner Agent provides strategic oversight, while specialized agents like the RAG Agent and the xBio-Tools Agent provide deep domain expertise in knowledge retrieval and computational modeling. This modular design mirrors the collaborative structure of human scientific teams and proves essential for navigating complex, multi-step research problems.

A key differentiator of BioLab is its ability to ground its reasoning in a rich, multimodal knowledge base and execute tasks using a suite of SOTA-level foundation models. The tripartite RAG system provides a dynamic and comprehensive view of biological knowledge, while the integration of the xTrimo Universe empowers BioLab to perform high-fidelity, quantitative predictions. These architectural features provide the foundation for the system’s advanced capabilities.

We first demonstrated these capabilities in a fully in silico case study, where BioLab autonomously executed an entire discovery workflow for a macrophage-targeting antibody. The system progressed from target identification and prioritization to the rational engineering of an optimized antibody candidate, a process that traditionally requires extensive human expertise and computational effort. This case study serves as a powerful demonstration of BioLab’s capacity for end-to-end automation within a virtual environment.

Building upon this, the most stringent validation of our platform was presented in the T cell study, where BioLab successfully closed the computational-experimental loop. This moved beyond pure simulation to guide physical experiments. The system not only autonomously identified PD-1 as the primary therapeutic target but also generated a detailed, actionable protocol that guided successful wet-lab knockout experiments. This ability to translate a computational strategy into a physical experimental plan represents a critical step towards true research automation. Moreover, BioLab’s subsequent optimization of the clinical antibody Pembrolizumab yielded novel variants with demonstrably superior potency in functional assays. This result prospectively validates the entire workflow, proving that BioLab can guide research toward tangible, high-value biological outcomes and significantly shorten the discovery-to-validation cycle.

Despite these advances, we acknowledge the limitations of the current BioLab system. Its ability to incorporate experimental feedback is currently dependent on human-mediated data entry, and the full automation of complex, physical wet-lab procedures remains a significant long-term challenge. The system’s knowledge is also constrained by the scope and quality of public databases and scientific literature. Future iterations will focus on developing more seamless interfaces for real-time experimental data ingestion and expanding the xTrimo Universe with models trained on proprietary, multi-modal datasets.

In conclusion, BioLab serves as a powerful prototype for the next generation of scientific discovery platforms. By unifying strategic reasoning, deep domain knowledge, and high-performance computing, it provides a scalable solution to the fragmentation of the research process. The principles demonstrated here—of modular agency, closed-loop learning, and the tight integration of computation and experimentation—lay the groundwork for a future where AI systems act not merely as tools, but as true collaborative partners in the quest for scientific understanding.

## Methods

### The BioLab multi-agent system

BioLab is implemented as a modular multi-agent system designed to automate closed-loop life science workflows, spanning in silico modeling and, where applicable, wet-lab experimentation. Drawing inspiration from cognitive architectures, BioLab organizes a collection of specialized agents into a structured hierarchy that enables dynamic task decomposition, inter-agent reasoning, model and tool invocation, and iterative refinement. Each agent is instantiated with a specific functional role, and agents interact via structured message-passing protocols that support both forward task progression and retrospective error correction. The core agents are detailed below.

#### Planner Agent

The system’s top-level orchestrator. Upon receiving a high-level scientific objective, the Planner Agent decomposes the task into a logical sequence of discrete subgoals. It then allocates these subtasks to the most appropriate downstream agents and manages the overall workflow to ensure the primary objective is met.

#### Reasoner Agent

The core of the system’s modular intelligence. The Reasoner Agent receives subgoals from the Planner Agent and is responsible for formulating detailed, scientifically valid action plans. It validates the internal coherence of proposed steps, assembles the required agent modules for execution, and coordinates the flow of information across upstream and downstream dependencies.

#### Memory Agent

This agent enables longitudinal learning and adaptive behavior. It maintains both short-term memory for the context of ongoing tasks and a long-term, versioned memory that stores execution traces, experimental outcomes, reusable agent configurations, and prior user feedback. This allows BioLab to reference past workflows and dynamically adjust its strategy based on empirical results.

#### RAG Agent

This agent grounds all system operations in factual evidence. It interfaces with a custom-built, multimodal knowledge base that includes structured databases (e.g., UniProt, PDB, GEO), semi-structured resources like protocol repositories, and the full text of the scientific literature. The RAG Agent is responsible for retrieving relevant information to support hypothesis generation, tool configuration, and experimental planning.

#### xBio-Tools Agent

This agent serves as the execution engine for a diverse collection of biological models and analytical tools. It abstracts the underlying software, including the xTrimo Universe foundation models (e.g., xTrimoSingleCellPerturb, xTrimoAntibody) and third-party tools, handling parameter translation, batch dispatch, and the return of structured, machine-readable results.

#### Code Agent

This suite of agents handles all code-related tasks. The Code Agent translates high-level agent workflows into executable Python or shell scripts. The Executor Agent then deploys these scripts in appropriate computational environments (e.g., containerized or HPC) and monitors their status. When failures or unexpected outputs arise, the Reflection Agent intervenes to perform root-cause analysis and propose remediation, such as revised code or model substitutions.

#### Critic Agent

The system’s quality control module. The Critic Agent continuously assesses the outputs of other agents against expected structural, statistical, or semantic criteria. It plays a central role in error detection and validation, triggering replanning or refinement cycles as necessary to ensure the scientific and technical rigor of the entire process.

#### Reporter Agent

The final agent in the workflow, responsible for communication. The Reporter Agent consolidates all data, analyses, and provenance metadata from a research cycle into comprehensive, human-readable reports. These reports are tailored for end-users with diverse backgrounds, from wet-lab biologists to computational modelers, ensuring the interpretability and auditability of the results.

The full agent system operates asynchronously with shared context layers and versioned memory objects, enabling reproducible, interpretable, and evolvable scientific execution. Throughout each cycle, BioLab agents collectively exhibit not only domain-specific capabilities but also adaptive system-level reasoning—a critical property for reliable, autonomous life science research.

### Memory Agent

The Memory Agent implements a hierarchical memory architecture for context retention and iterative improvement. It comprises a Short-Term Memory (STM) that maintains transient, task-level context and a Long-Term Memory (LTM) that stores and retrieves persistent knowledge. The design follows standard agentic-memory practice and, where noted, draws on ideas from the mem0^61^ framework.

#### Short-term memory

STM operates as a high-speed “working memory”, maintaining a rolling window of structured records produced during an active workflow. Each record is timestamped and annotated with its source agent and artifact type (e.g., tool output, intermediate reasoning). The buffer is fixed-length with first-in–first-out eviction; agents query it during execution to ensure coherence across multi-step procedures. On task completion, a summarization routine condenses the session log and emits a compact representation of salient facts and decisions for downstream consolidation.

#### Long-term memory

LTM serves as the persistent knowledge base and is organized into two stores. A user-profile store (key–value) maintains user-specific parameters, project history, and tool configurations keyed by a unique user id, enabling context-aware behavior across sessions. A factual store (vector) maintains validated facts, procedural workflows, and experimental findings as high-dimensional embeddings; similarity search over this store supports semantic retrieval and analogical reuse across problems. Retrieval uses standard vector similarity with approximate nearest-neighbour indexing; store updates are transactionally logged to preserve provenance.

#### Consolidation and update

After each task, an LLM-driven consolidation pass extracts candidate memories from the STM summary, queries the factual store for the top-*k* nearest neighbours (configurable), and compares candidates with retrieved entries. The model issues one of four operations—ADD (novel entry), UPDATE (refinement or correction), DELETE (retire obsolete entry), or NOOP (no change)—and writes an audit record containing the pre/post states and rationale. User feedback on retrieval quality and tool performance is ingested as factual entries and subsequently influences retrieval weighting and tool-specific confidence scores. This closed loop yields gradual improvement in retrieval precision and downstream decision quality without altering task-time behaviour.

### RAG Agent and Multi-modal Knowledge Base

To ground all reasoning in robust, verifiable scientific evidence and mitigate the risk of factual hallucination, BioLab incorporates a highly specialized Retrieval-Augmented Generation (RAG) Agent. This agent functions as an advanced information retrieval and synthesis engine, interfacing between the reasoning core of BioLab and a custom-built, multimodal knowledge base. The RAG Agent employs a multi-stage pipeline that includes query pre-processing, parallel hybrid retrieval across diverse knowledge sources, and intelligent information fusion to provide a rich, contextually relevant evidence base for downstream tasks. Detailed information about RAG Agent and the multi-modal knowledge base is in Extended Data Figure S1.

#### Query Pre-processing

Upon receiving a query from an upstream agent (e.g., the Reasoner), the RAG Agent first initiates a query pre-processing step. A dedicated Query Rewrite LLM Agent analyzes the initial query and systematically expands it into a set of comprehensive sub-queries. This expansion is performed across multiple dimensions to maximize retrieval coverage, including: (i) synonym and related term expansion to capture lexical variations; (ii) thematic expansion with related biological concepts; (iii) contextual expansion based on the preceding conversational history; and (iv) multi-angle expansion, where the query is rephrased to seek definitions, causes, consequences, etc. This process transforms a single, often ambiguous, query into a rich set of specific questions poised for deep retrieval.

#### Hybrid Retrieval from a Multimodal Knowledge Base

The generated sub-queries are then dispatched in parallel to three specialized retrieval channels, each targeting a distinct component of our multimodal knowledge base. This base was designed to provide a comprehensive and synergistic view of biomedical knowledge.

- **Document Search**: This channel targets a curated library of scientific literature and internal documents. It employs a hybrid retrieval strategy, combining dense (vector-based semantic search) and sparse (e.g., BM25 keyword-based) retrieval methods to ensure both conceptual and lexical relevance. The results are further refined using metadata filters.
- **Web Search**: To access the most current information, a Web Search channel generates optimized search engine queries from the sub-queries. It retrieves and processes content from a variety of web sources, including HTML pages and PDF documents, using specialized document loaders.
- **Knowledge Graph Search**: This channel provides access to structured, factual knowledge. An LLM first performs named entity recognition on the sub-queries to identify key biological entities. These entities are then mapped to nodes in our proprietary biomedical knowledge graph. A graph query (e.g., Cypher) is then generated to traverse the graph and retrieve precise, factual relationships between entities. The architecture of this knowledge base is detailed in Extended Data Figure S1.

#### Information Fusion and Reranking

The raw outputs from the three parallel retrieval channels are not used directly. Instead, they are fed into a sophisticated fusion and reranking module. A DeepSource Master LLM Agent first assesses the relevance and quality of the retrieved information chunks from each source. Concurrently, a Global Multi-Channel Weighting model assigns a confidence score to each chunk based on its source, relevance, and timeliness. The top-’k’ ranked information snippets, weighted by their scores, are then synthesized and consolidated into a final, coherent context block. This reranking and fusion step is critical for filtering out noise and ensuring that only the highest-quality, most relevant evidence is passed to the final generation stage. This refined context is then provided to the downstream agent that initiated the query, enabling it to generate a response that is deeply informed and factually grounded.

### Benchmarking and evaluation methodology

To systematically evaluate the performance of BioLab’s reasoning capabilities, we curated a comprehensive suite of benchmarks spanning biomedical question answering, multi-hop scientific reasoning, and open-ended research task planning. These benchmarks were selected to test not only factual recall, but also the ability to integrate domain knowledge, synthesize context, and generate expert-level research strategies. We compare BioLab against state-of-the-art large language models (LLMs) and agent-based baselines using both quantitative metrics and expert evaluation.

#### Biomedical reasoning benchmarks

We first evaluated BioLab on a collection of four domain-specific reasoning benchmarks. These included PubMedQA (both with and without contextual passage)^34^, MMLU-Pro-Biology (with chain-of-thought prompting)^35^, and GPQA-Diamond^36^. PubMedQA is a question-answering benchmark designed to assess biomedical knowledge and reasoning using questions sourced from PubMed abstracts. It consists of 1,000 expert-annotated yes/no/maybe questions, each paired with a relevant abstract in the contextual passage setting, requiring models to extract and reason over pertinent information. In the no-context setting, models must rely solely on prior knowledge to answer questions, testing their ability to recall and apply biomedical facts without external references. This dual setup evaluates both information retrieval and standalone reasoning capabilities. MMLU-Pro-Biology, a subset of the Massive Multitask Language Understanding (MMLU) benchmark, focuses on advanced biology questions at the college and professional level. It comprises 1,200 multiple-choice questions covering topics such as molecular biology, genetics, and physiology. With chain-of-thought prompting, models are encouraged to generate step-by-step reasoning before selecting an answer, emphasizing their ability to perform complex, multi-step reasoning in biology. This benchmark tests deep conceptual understanding and the capacity to connect disparate biological concepts. GPQA-Diamond is a highly curated benchmark of 198 graduate-level biology questions, designed to evaluate grounded reasoning in specialized biomedical domains. Each question is crafted by domain experts to require both factual knowledge and sophisticated reasoning, often involving the integration of multiple scientific concepts. The benchmark emphasizes real-world applicability, with questions reflecting challenges encountered in cutting-edge biomedical research. Performance is measured in percentiles, highlighting a model’s ability to excel in expert-level reasoning tasks.

#### Expert-graded research task planning benchmark

To evaluate BioLab’s planner and reasoning agents in open-ended contexts, we constructed a custom benchmark of 20 research questions named BioResearchQA across molecular biology, immunology, RNA therapeutics, protein design, and synthetic cell engineering. Each question was formulated as a realistic research problem—for instance, “Design a strategy to identify non-coding RNA regulators of macrophage polarization”—requiring multi-step planning and domain-specific tool selection. Three domain experts scored each output along three axes: (1) multi-modal planning, (2) bio-tool calling, and (3) full-loop science reasoning. Scoring used a 5-point Likert scale per criterion, averaged across reviewers.

#### LLM baselines

To contextualize BioLab’s agentic reasoning capabilities, we conducted additional evaluations against leading LLMs including DeepSeek-R1^37^, Qwen3-235B-A22B^38^, Gemini-2.5-Flash^41^, Gemini-2.5-Pro^42^, and ChatGPT-4o^39, 40^. DeepSeek-R1, developed by DeepSeek AI, is a 671B-parameter dense model optimized for reasoning and coding, with a 131,072-token context window and strong performance on benchmarks^37^. Qwen3-235B-A22B, from Alibaba’s Qwen team, employs a Mixture-of-Experts (MoE) architecture with 235B total parameters (22B active), excelling in coding (LiveCodeBench: 70.7) and math (AIME 2024: 85.7) under the Apache 2.0 license^38^. Gemini-2.5-Flash, a lightweight model from Google, prioritizes low-latency tasks^41^, while Gemini-2.5-Pro, a more robust counterpart, leads in reasoning benchmarks like ArenaHard (96.4%)^42^. ChatGPT-4o, developed by OpenAI, is a multimodal model with strong general-purpose capabilities, though it slightly trails Qwen3-235B-A22B in coding tasks like Codeforces (2056 Elo vs. 1974 Elo)^39, 40^. These models represent a spectrum of architectural and deployment strategies, evaluated here for their biomedical reasoning capabilities.

#### Agent-based systems

This study examines an advanced AI agents designed for specialized reasoning and task execution: Biomni^62^. Biomni is a general-purpose biomedical AI agent that autonomously executes tasks across 25 biomedical subfields. By combining a unified action space (Biomni-E1) with a dynamic task-executing architecture (Biomni-A1), it handles complex workflows like multi-omics analysis, achieving 74.4% accuracy on LAB-Bench DbQA and 81.9% on SeqQA, surpassing human expert performance^62^. The agent-based system represent cutting-edge approaches to autonomous reasoning, tool integration, and domain-specific problem-solving in AI research.

### xTrimo platform integration and foundation model usage within BioLab

The BioLab system integrates a suite of domain-specialized foundation models through its modular xTrimo platform, which serves as a unified inference backbone for biological sequence modeling, perturbation prediction, and structural optimization. The xTrimo platform includes both general-purpose models for protein, RNA, and DNA analysis, as well as task-specific foundation models designed for high-fidelity interpretation of cellular and molecular function. These models are invoked by BioLab agents via the xBio-Tools interface, which handles model selection, parameter configuration, and input/output formatting for downstream reasoning.

Among the core components of this platform is xTrimoSingleCellPerturb based on xTrimoCell. xTrimoCell is a single-cell foundation model purpose-built for predicting genotype-to-phenotype responses to genetic perturbations. Trained on over 5 million single-cell expression profiles and fine-tuned with a large-scale perturb-seq dataset comprising 347,738 cells, xTrimoCell captures context-dependent transcriptional responses across diverse cell types. The model architecture leverages multi-head attention and contrastive alignment across cell embeddings and perturbation graphs, enabling robust generalization to previously unseen gene–cell–condition triplets.

In BioLab, xTrimoSingleCellPerturb is accessed by the Reasoner and Target Identification agents during hypothesis formation and candidate ranking. For example, when the system is tasked with nominating modulators of T cell exhaustion or macrophage activation, it queries xTrimoSingleCellPerturb with context-specific perturbation sets. The model outputs predicted transcriptional shifts, which are ranked according to inferred immune activation markers (e.g., IFNG, GZMB) and integrated with graph-based literature mining results via the Retriever Agent. These integrated scores are used by BioLab to prioritize gene targets for wet-lab validation or therapeutic development.

Complementing this is xTrimoAntibody based on xTrimoPGLM^63^. xTrimoPGLM is a multi-billion-parameter biomolecular foundation model designed for unified sequence–structure reasoning across proteins and other biopolymers. xTrimoPGLM incorporates autoregressive and masked modeling heads, enabling it to support both generative and discriminative tasks such as affinity scoring, structural refinement, and variant prediction. The model is pre-trained on over 1.2 billion natural and synthetic sequences, with co-training over known structures and fine-tuning on experimental protein–ligand interaction data.

Within BioLab, the xTrimoAntibody model is called by the Structure Reasoning and Optimization agents to evaluate candidate therapeutic designs. In antibody development tasks, for instance, it is used to assess interface complementarity between antibody–antigen pairs, to compute developability metrics including aggregation risk and thermostability, and to guide multi-objective optimization routines. The model’s generative capacity is leveraged during sequence refinement, where candidate antibodies are iteratively modified and scored to balance binding affinity, structural validity, and sequence novelty.

By enabling task-specific, context-aware modeling at both cellular and molecular scales, the integration of xTrimoSingleCellPerturb and xTrimoPGLM allows BioLab to reason over biological systems with a degree of precision and adaptability not achievable using generic LLMs alone. The models are further extensible within the BioLab platform, supporting modular composition with downstream predictors and facilitating robust generalization to novel experimental settings.

### Molecule Dynamics Simulation and Analysis of Wild-Type and Optimized Antibody-Antigen

#### Complexes System Preparation and Simulation Parameters

To investigate the structural basis of antibody–antigen recognition, all-atom MD simulations were performed for the wild-type and optimized antibody–receptor complexes. Initial structures were prepared and subsequently simulated using GROMACS (version 2021.4) with the CHARMM36m force field, which is well-validated for protein dynamics. Each complex was solvated in a dodecahedron water box using the TIP3P water model, ensuring a minimum distance of 10 °A between the protein and the box edges to prevent periodic image artifacts. The system was neutralized and brought to a physiological salt concentration of 0.15 M by adding Na^+^ and Cl^−^ counter-ions. Long-range electrostatic interactions were computed using the Particle Mesh Ewald (PME) method with a real-space cutoff of 12 °A. A corresponding 12 °A cutoff was applied for Lennard–Jones interactions, with a force-based switching function initiated at 10 °A.

#### System Equilibration and Production Simulations

Prior to production runs, each system underwent a rigorous multi-step equilibration protocol to ensure thermal and structural stability. First, an energy minimization was performed using the steepest descent algorithm to remove any steric clashes. The system was then gradually heated to 300 K over 1 ns in the canonical (NVT) ensemble, with position restraints applied to all protein heavy atoms to allow the solvent to relax. This was followed by a 5 ns equilibration in the isothermal-isobaric (NPT) ensemble at 1 atm, using an isotropic Parrinello–Rahman barostat, to ensure proper system density. During this phase, the position restraints on the heavy atoms were progressively released. All covalent bonds involving hydrogen atoms were constrained using the LINCS algorithm, which enabled a stable 2 fs integration time step. For robust sampling, three independent production simulations, each starting from different initial velocities, were carried out for 100 ns per system.

#### Conformational Clustering and Trajectory Analysis

To identify the most representative and thermodynamically stable protein conformations from the dynamic trajectories, all post-simulation analyses were performed on the concatenated trajectories from the three independent replicates for each system. Conformational clustering was performed using the gmx cluster tool in GROMACS with the Gromos algorithm. A 1.5 °A root-mean-square deviation (RMSD) cutoff, calculated on the backbone atoms of the interface residues (defined as residues with any atom within 10 °A of the binding partner), was used to group similar structures. The centroid structure from the most populated cluster for each system, defined as the frame with the smallest average RMSD to all other frames in that cluster, was selected for all subsequent detailed analyses.

#### Interfacial Fingerprinting and Visualization

The interfacial fingerprint for each antibody complex was constructed by computing and mapping key physicochemical properties onto its molecular surface. Starting from the selected representative centroid structure, a triangulated surface mesh was generated at 1.0 °A resolution using MSMS. Interfacial vertices were defined as surface points on the antibody within 4 °A of any receptor atom. For each interfacial vertex, four distinct feature channels were calculated: (i) Shape index, to describe the local surface curvature (concave, convex, or flat); (ii) Poisson–Boltzmann electrostatic potential, calculated using the APBS software to map charge distribution; (iii) Hydropathy, based on the Kyte-Doolittle scale to represent hydrophobicity; and (iv) Hydrogen-bonding potential, which includes separate features for lone-pair electron acceptors and proton donors to map binding capability. Each feature channel was normalized to a range of [0, 1] and projected onto the antibody interface. All structural visualizations were generated using PyMOL.

### Wet-lab validation of T cell cycle predictions and antibody candidates

To validate BioLab’s computational predictions in a real-world biological context, we conducted wet-lab experiments focused on T cell functional modulation and therapeutic antibody optimization. These experiments were performed in collaboration with a certified immunology laboratory and followed standardized experimental procedures for genome perturbation and antibody characterization.

#### CRISPR perturbation of candidate T cell targets

Following computational nomination of immune checkpoint targets by BioLab’s xTrimoSingleCellPerturb-based inference, CRISPR–Cas9–mediated knockdown experiments were designed using sgRNA sequences automatically generated by BioLab’s sequence design agent. The sgRNAs were selected based on predicted on-target efficacy and off-target avoidance, incorporating integrated rules from *CRISPRscan* and Doench (2016) scoring. Primary human CD8^+^ T cells were isolated from peripheral blood mononuclear cells (PBMCs) via negative selection and cultured in the presence of IL-2 (100 IU/mL). Nucleofection was performed using a Lonza 4D-Nucleofector (program EO-115) with pre-complexed sgRNA/Cas9 ribonucleoproteins. Cells were harvested 72 hours post-transfection.

Functional outcomes were assessed using flow cytometry and cytokine profiling. Cells were stained for activation markers (CD69, CD25) and intracellular IFN-*γ* and Granzyme B. Data acquisition was performed on a BD FACSCelesta flow cytometer, and analyses were conducted in FlowJo v10. Statistical comparisons were performed using one-way ANOVA followed by Dunnett’s post hoc test for multiple comparisons, with adjusted *p*-values *<* 0.05 considered significant.

#### Antibody expression and purification

Candidate antibody sequences optimized by BioLab’s xTrimoAntibody agent were synthesized and cloned into IgG1 heavy- and light-chain expression vectors. Constructs were transiently transfected into Expi293F cells using ExpiFectamine (Thermo Fisher) and cultured in Expi293 Expression Medium at 37°C, 8% CO_2_, shaking at 125 rpm. Supernatants were harvested at day 7 post-transfection and purified using Protein A affinity chromatography. Antibody concentration was quantified by absorbance at 280 nm and validated by SDS–PAGE.

#### Binding and functional assays

Binding affinity to PD-1 was measured using bio-layer interferometry (BLI) on an Octet RH96 system. Anti-human IgG Fc capture (AHC) biosensors were loaded with antibody samples (5 *μ*g/mL), and association/dissociation kinetics were recorded against recombinant human PD-1–Fc (R&D Systems). Equilibrium dissociation constants (*K*_*D*_) were determined using a 1:1 binding model in the ForteBio Data Analysis software. Functional potency was assessed using a PD-1 reporter assay. Jurkat–NFAT–P3Z cells were co-cultured with CHO–K1–PD-L1 cells in the presence of serially diluted antibody samples. Luciferase activity was measured after 6 h using the *Bright-Glo* Luciferase Assay (Promega), and IC_50_ values were calculated by four-parameter logistic regression in GraphPad Prism v9.

#### Stability characterization

Thermal stability was assessed using nano differential scanning fluorimetry (NanoDSF, Panta Nano DSF). Protein samples were centrifuged at 3000 × g for 10 min, and the supernatant was loaded into glass capillaries. Samples were scanned from 25 °C to 95 °C at a heating rate of 1 °C / min. The melting temperature (*T*_*m*_) was determined as the inflection point of the unfolding transition using the Panta Analysis software, with reference controls confirming system suitability (*T*_*m*1_ within 0.3 ±°C).

#### Hydrophobic interaction chromatography (HIC) analysis

HIC was performed to assess protein retention time and aggregation potential, using a TSKgel Butyl-NPR column (Tosoh Bioscience) on an Agilent HPLC system with ammonium sulfate gradients.

### Statistical Analysis

Unless otherwise stated, all assays were performed in at least three biological replicates. Data are presented as mean ± s.e.m. (standard error of the mean). Significance tests were two-sided unless otherwise specified, and a *p*-values *<* 0.05 was considered statistically significant. For comparisons involving multiple groups or conditions, one-way or two-way ANOVA was used, followed by an appropriate post hoc test for multiple comparisons (e.g., Dunnett’s or Bonferroni’s correction). All statistical analyses were conducted in R (v4.2.1) or GraphPad Prism (v9).

## Computational Resources

All computational experiments—including agent execution, model inference, and optimization workflows—were performed on a high-performance computing cluster equipped with NVIDIA A100 GPUs (40GB VRAM) and 1.5 TB of aggregate system memory. Foundation model inference was conducted using containerized services deployed via Docker, orchestrated through a custom task manager built on the AutoGen framework. Training and evaluation pipelines were implemented in Python 3.10 with PyTorch 2.1 and Hugging Face Transformers 4.38, and executed on Ubuntu 22.04 LTS. Agent coordination followed an event-driven architecture using asynchronous message-passing, with real-time state tracking via Redis-backed memory. Multi-agent episodes were logged and versioned using MLflow to ensure reproducibility across runs. All experiments were executed under deterministic seed initialization unless otherwise noted.

## Code availability

The source code for the BioLab multi-agent system and the models used in this study will be made publicly available on a dedicated GitHub repository upon publication of this manuscript. In the interim, the core functionalities of BioLab are available for exploration via an interactive web-based platform at https://xtrimo.biomap.com/.

## Author contributions statement

R.J., R.Z., Z.Z., M.W., X.Z., and L.C. conceived the study and designed the experiments. R.J., Y.G., Y.Q., and M.Y. performed the dataset analyses and developed the foundation models. R.J., C.S., Q.Y., L.L.C., Y.Z., R.X., and Z.X. conducted the downstream experiments and collected the data. R.J., Y.G., and Y.Q. analyzed the data and interpreted the results. R.Z., Z.Z., M.W., X.Z., and L.C. supervised the project and provided critical intellectual input. R.J., Y.G., and Y.Q. wrote the initial draft of the manuscript. Y.Z., R.X., and Z.X. assisted with manuscript editing. R.Z., Z.Z., M.W., X.Z., and L.C. revised the manuscript. All authors reviewed and approved the final version of the manuscript.

## Additional information

**Correspondence and requests for materials** should be addressed to Ruhong Zhou, Zaixi Zhang, Mengdi Wang, Xiaoming Zhang, and Le Cong.

## References

1. Rozenblatt-Rosen, O., Stubbington, M. J. T., Regev, A. & Teichmann, S. A. The human cell atlas: from vision to reality. Nature 550, 451–453 (2017). URL 10.1038/550451a.

2. Tabula Sapiens Consortium. The tabula sapiens: A multiple-organ, single-cell transcriptomic atlas of humans. Science 376, eabl4896 (2022). URL 10.1126/science.abl4896.

3. Lai, Y. et al. Multimodal cell atlas of the ageing human skeletal muscle. Nature 630, 345–352 (2024). URL 10.1038/s41586-024-07715-3.

4. Backman, J. D. et al. Whole-genome sequencing of 490,640 uk biobank participants. Nature 622, 472–480 (2023). URL 10.1038/s41586-025-09272-9.

5. Canty, R. B. et al. Science acceleration and accessibility with self-driving labs. Nature Communications 16, 301 (2025). URL 10.1038/s41467-025-59231-1.

6. Liu, T. et al. Evaluating the utilities of foundation models in single-cell data analysis. bioRxiv (2023). URL 10.1101/2023.09.08.555192.

7. Uzzi, B., Mukherjee, S., Stringer, M. & Jones, B. Atypical combinations and scientific impact. Science 342, 468–472 (2013). URL https://www.science.org/doi/10.1126/science.1240474. https://www.science.org/doi/pdf/10.1126/science.1240474.

8. Wilkinson, M. D., Dumontier, M., Aalbersberg, I. J. et al. The fair guiding principles for scientific data management and stewardship. Scientific Data 3, 160018 (2016). URL 10.1038/sdata.2016.18.

9. Szymanski, N. J. et al. An autonomous laboratory for accelerated synthesis of inorganic materials. Nature 620, 66–73 (2023). URL 10.1038/s41586-023-06734-w.

10. Rapp, J. T. et al. Self-driving laboratories to autonomously navigate the protein fitness landscape. Nature 624, 64–71 (2024). URL 10.1038/s44286-023-00002-4.

11. Scannell, J. W., Blanckley, A., Boldon, H. & Warrington, B. Diagnosing the decline in pharmaceutical r&d efficiency. Nature Reviews Drug Discovery 11, 191–200 (2012). URL 10.1038/nrd3681.

12. Ma, Q. et al. Harnessing the deep learning power of foundation models in single-cell omics. Nature Reviews Molecular Cell Biology 25, 5–22 (2024). URL 10.1038/s41580-024-00756-6.

13. Ahlmann-Eltze, C. et al. Deep-learning-based gene perturbation effect prediction does not yet outperform simple linear baselines. Nature Methods (2025). URL 10.1038/s41592-025-02772-6.

14. Rives, A. et al. Biological structure and function emerge from scaling unsupervised learning to 250 million protein sequences. Proceedings of the National Academy of Sciences 118, e2016239118 (2021). URL 10.1073/pnas.2016239118. https://www.pnas.org/doi/pdf/10.1073/pnas.2016239118.

15. Xu, F., Wu, T., Cheng, Q., Wang, X. & Yan, J. Foundation models in plant molecular biology: advances, challenges, and future directions. Frontiers in Plant Science 16, 1611992 (2025). URL 10.3389/fpls.2025.1611992.

16. Nguyen, E. et al. Sequence modeling and design from molecular to genome scale with evo. Science 386, eado9336 (2024). URL https://www.science.org/doi/10.1126/science.ado9336. https://www.science.org/doi/pdf/10.1126/science.ado9336.

17. Boiko, D. A., MacKnight, R. & Gomes, G. Emergent autonomous scientific research capabilities of large language models. arXiv preprint 2304.05332 (2023). URL 10.48550/arXiv.2304.05332.

18. Ball, P. Is ai leading to a reproducibility crisis in science? Nature 624, 22–25 (2023). URL 10.1038/d41586-023-03817-6.

19. Kitano, H. Nobel turing challenge: creating the engine for scientific discovery. npj Systems Biology and Applications 7, 29 (2021). URL 10.1038/s41540-021-00189-3.

20. Canty, R. B. et al. Science acceleration and accessibility with self-driving labs. Nature Communications 16, 3856 (2025). URL 10.1038/s41467-025-59231-1.

21. Boiko, D. A., MacKnight, R., Kline, B. & Gomes, G. Autonomous chemical research with large language models. Nature 624, 570–578 (2023). URL 10.1038/s41586-023-06792-0.

22. Horawalavithana, S. et al. Foundation models of scientific knowledge for chemistry: Opportunities, challenges and lessons learned. In Fan, A., Ilic, S., Wolf, T. & Gallé, M. (eds.) Proceedings of BigScience Episode #5 – Workshop on Challenges & Perspectives in Creating Large Language Models, 160–172 (Association for Computational Linguistics, virtual+Dublin, 2022). URL https://aclanthology.org/2022.bigscience-1.12/.

23. Chen, H. et al. An overview of domain-specific foundation model: key technologies, applications and challenges. arXiv preprint 2409.04267 (2024). URL 10.48550/arXiv.2409.04267.

24. Zhou, C., Li, Q., Li, C. et al. A comprehensive survey on pretrained foundation models: a history from bert to chatgpt. International Journal of Machine Learning and Cybernetics (2024). URL 10.1007/s13042-024-02443-6.

25. Singhal, K., Azizi, S., Tu, T. et al. Large language models encode clinical knowledge. Nature 620, 172–180 (2023). URL 10.1038/s41586-023-06291-2.

26. Qian, C., Tang, H., Yang, Z., Liang, H. & Liu, Y. Can large language models empower molecular property prediction? arXiv preprint 2307.07443 (2023). URL 10.48550/arXiv.2307.07443.

27. Nori, H. et al. Can generalist foundation models outcompete special-purpose tuning? case study in medicine. arXiv preprint 2311.16452 (2023). URL 10.48550/arXiv.2311.16452.

28. Thirunavukarasu, A. J., Ting, D. S. J., Elangovan, K. et al. Large language models in medicine. Nature Medicine 29, 1930–1940 (2023). URL 10.1038/s41591-023-02448-8.

29. Schölkopf, B. et al. Towards causal representation learning. arXiv preprint 2102.11107 (2021). URL https://doi.org/10.48550/arXiv.2102.11107.

30. Huang, L. et al. A survey on hallucination in large language models: Principles, taxonomy, challenges, and open questions. ACM Trans. Inf. Syst. 43 (2025). URL 10.1145/3703155.

31. Ji, Z. et al. Survey of hallucination in natural language generation. ACM Comput. Surv. 55, 248:1–248:38 (2023).

32. Seyhan, A. A. Lost in translation: the valley of death across preclinical and clinical divide – identification of problems and overcoming obstacles. Translational Medicine Communications 4, 18 (2019). URL 10.1186/s41231-019-0050-7.

33. Macklin, P. Key challenges facing data-driven multicellular systems biology. arXiv preprint 1806.04736 (2018). URL https://arxiv.org/abs/1806.04736. Version 2, last revised 27 Sep 2019.

34. Jin, Q., Dhingra, B., Liu, Z., Cohen, W. W. & Lu, X. Pubmedqa: A dataset for biomedical research question answering. arXiv preprint 1909.06146 (2019). URL 10.48550/arXiv.1909.06146.

35. Wang, Y. et al. Mmlu-pro: A more robust and challenging multi-task language understanding benchmark. arXiv preprint 2406.01574 (2024). URL 10.48550/arXiv.2406.01574.

36. Rein, D. et al. Gpqa: A graduate-level google-proof q&a benchmark. arXiv preprint 2311.12022 (2023). URL 10.48550/arXiv.2311.12022.

37. DeepSeek-AI et al. Deepseek-r1: Incentivizing reasoning capability in llms via reinforcement learning. arXiv preprint 2501.12948 (2025). URL https://arxiv.org/abs/2501.12948.

38. Yang, A. et al. Qwen3 technical report. arXiv preprint 2505.09388 (2025). URL https://arxiv.org/abs/2505.09388.

39. OpenAI. Hello gpt-4o. https://openai.com/index/hello-gpt-4o/(2024). Accessed: 2025-08-15.

40. OpenAI et al. Gpt-4o system card. arXiv preprint 2410.21276 (2024). URL https://arxiv.org/abs/2410.21276.

41. DeepMind, G. Gemini 2.5 flash – fast performance, native multimodality, and long-context capability. https://deepmind.google/models/gemini/flash/(2025). Accessed: 2025-08-15.

42. Comanici, G. et al. Gemini 2.5: Pushing the frontier with advanced reasoning, multimodality, long context, and next generation agentic capabilities. arXiv preprint 2507.06261 (2025). URL 10.48550/arXiv.2507.06261.

43. Zhang, S. et al. Position: Intelligent science laboratory requires the integration of cognitive and embodied ai. arXiv preprint 2506.19613 (2025). URL 10.48550/arXiv.2506.19613.

44. Ono, T., Hayashi, M., Sasaki, F. & Nakashima, T. Rankl biology: bone metabolism, the immune system, and beyond. Inflammation and Regeneration 40, 2 (2020). URL 10.1186/s41232-019-0111-3.

45. Anderson, D., Maraskovsky, E., Billingsley, W. et al. A homologue of the tnf receptor and its ligand enhance t-cell growth and dendritic-cell function. Nature 390, 175–179 (1997). URL 10.1038/36593.

46. Lacey, D. L. et al. Osteoprotegerin ligand is a cytokine that regulates osteoclast differentiation and activation. Cell 93, 165–176 (1998). URL 10.1016/S0092-8674(00)81569-X.

47. Yasuda, H. et al. Osteoclast differentiation factor is a ligand for osteoprotegerin/osteoclastogenesis-inhibitory factor and is identical to trance/rankl. Proceedings of the National Academy of Sciences of the United States of America 95, 3597–3602 (1998). URL 10.1073/pnas.95.7.3597.

48. Hanley, D. A., Adachi, J. D., Bell, A. & Brown, V. Denosumab: mechanism of action and clinical outcomes. International Journal of Clinical Practice 66, 1139–1146 (2012). URL 10.1111/ijcp.12022. Epub 2012 Sep 12; PubMed PMID: 22967310; PMCID: PMC3549483.

49. Cummings, S. R. et al. Denosumab for prevention of fractures in postmenopausal women with osteoporosis. New England Journal of Medicine 361, 756–765 (2009). URL 10.1056/NEJMoa0809493.

50. Kostenuik, P. J. et al. Denosumab, a fully human monoclonal antibody to rankl, inhibits bone resorption and increases bmd in knock-in mice that express chimeric (murine/human) rankl. Journal of Bone and Mineral Research 24, 182–195 (2009). URL 10.1359/jbmr.081112.

51. Thomas, D. et al. Denosumab in patients with giant-cell tumour of bone: an open-label, phase 2 study. The Lancet Oncology 11, 275–280 (2010). URL 10.1016/S1470-2045(10)70010-3. Epub 2010 Feb 10.

52. Alder, B. J. & Wainwright, T. E. Studies in molecular dynamics. i. general method. The Journal of Chemical Physics 31, 459–466 (1959).

53. McCammon, J. A., Gelin, B. R. & Karplus, M. Dynamics of folded proteins. Nature 267, 585–590 (1977).

54. Jinek, M. et al. A programmable dual-rna–guided dna endonuclease in adaptive bacterial immunity. Science 337, 816–821 (2012).

55. Cong, L. et al. Multiplex genome engineering using crispr/cas systems. Science 339, 819–823 (2013).

56. Mali, P. et al. Rna-guided human genome engineering via cas9. Science 339, 823–826 (2013).

57. Groff-Vindman, C. S. et al. The convergence of ai and synthetic biology: the looming revolution. Nature (2025).

58. Ye, F. et al. Proteinbench: A holistic evaluation of protein foundation models. arXiv preprint 2409.06744 (2024). URL https://arxiv.org/abs/2409.06744.

59. Darwish, A. M., Rashed, E. A. & Khoriba, G. Mitigating llm hallucinations using a multi-agent framework. Information 16, 517 (2025). URL https://www.mdpi.com/2078-2489/16/7/517.

60. Kim, Y. et al. Medical hallucinations in foundation models and their impact on healthcare. arXiv preprint 2503.05777(2025). URL https://arxiv.org/abs/2503.05777.

61. Chhikara, P., Khant, D., Aryan, S., Singh, T. & Yadav, D. Mem0: Building production-ready ai agents with scalable long-term memory. arXiv preprint 2504.19413 (2025). URL https://arxiv.org/abs/2504.19413.

62. Huang, K. et al. Biomni: A general-purpose biomedical ai agent. bioRxiv (2025). URL https://www.biorxiv.org/content/early/2025/05/30/2025.05.30.656746.

63. Chen, B., Cheng, X., Li, P. et al. xtrimopglm: unified 100-billion-parameter pretrained transformer for deciphering the language of proteins. Nature Methods 22, 1028–1039 (2025). URL 10.1038/s41592-025-02636-z.

64. Yang, A. et al. Qwen2.5: A party of foundation models. arXiv preprint 2407.10671 (2024). URL https://qwenlm.github.io/blog/qwen2.5/.

65. Schulman, J., Wolski, F., Dhariwal, P., Radford, A. & Klimov, O. Proximal policy optimization algorithms. arXiv preprint 1707.06347 (2017). URL https://arxiv.org/abs/1707.06347.

66. Yang, Q. et al. Trinitydna: A bio-inspired foundational model for efficient long-sequence dna modeling. arXiv preprint 2507.19229 (2025). URL https://arxiv.org/abs/2507.19229. Version 1, submitted 25 Jul 2025.

67. Zhang, Z. et al. Rnagenesis: A generalist foundation model for functional rna therapeutics. bioRxiv (2025). URL https://www.biorxiv.org/content/early/2025/07/01/2024.12.30.630826.

68. Hao, M., Gong, J., Zeng, X. et al. Large-scale foundation model on single-cell transcriptomics. Nature Methods 21, 1481–1491 (2024). URL 10.1038/s41592-024-02305-7.

69. Hu, Z. et al. Drug synergistic combinations predictions via large-scale pre-training and graph structure learning. arXiv preprint 2301.05931 (2023). URL 10.48550/arXiv.2301.05931.

